# Interpersonal early adversity demonstrates dissimilarity from early socioeconomic disadvantage in the course of human brain development: A meta-analysis

**DOI:** 10.1101/2023.02.16.528877

**Authors:** Anna Vannucci, Andrea Fields, Eleanor Hansen, Ariel Katz, John Kerwin, Ayumi Tachida, Nathan Martin, Nim Tottenham

## Abstract

It has been established that early-life adversity impacts brain development, but the role of development itself has largely been ignored. We take a developmentally-sensitive approach to examine the neurodevelopmental sequelae of early adversity in a preregistered meta-analysis of 27,234 youth (birth to 18-years-old), providing the largest group of adversity-exposed youth to date. Findings demonstrate that early-life adversity does not have an ontogenetically uniform impact on brain volumes, but instead exhibits age-, experience-, and region-specific associations. Relative to non-exposed comparisons, interpersonal early adversity (e.g., family-based maltreatment) was associated with initially larger volumes in frontolimbic regions until ~10-years-old, after which these exposures were linked to increasingly smaller volumes. By contrast, socioeconomic disadvantage (e.g., poverty) was associated with smaller volumes in temporal-limbic regions in childhood, which were attenuated at older ages. These findings advance ongoing debates regarding why, when, and how early-life adversity shapes later neural outcomes.

## 1. Introduction

Exposure to early adverse experiences (caregiver separation, abuse, neglect, violence exposure, poverty, etc.), has outsized associations with the developing brain (Tierney and Nelson, 2009). A growing body of empirical work has demonstrated consistent links between early-life adversity exposure and structural maturation (Lim et al., 2018; McLaughlin et al., 2019; Tottenham, 2020). However, there is a large gap in this literature with regard to considering the role of development—that is, associations between early experiences and neural phenotypes are not developmentally monolithic and outcomes diverge as a function of both the nature of the adversity (Ford et al., 2011) and developmental timing (VanTieghem et al., 2021). It is therefore misleading to assume that brain outcomes linked to an early adverse experience measured during one developmental window (e.g., childhood) will be the same as those measured during another (e.g., adulthood). Here, we demonstrate the imperative of considering both adversity type and development to characterize the impact of early adverse experiences on brain structure. Importantly, successfully disentangling the complex associations between early adverse experiences and neurodevelopment requires large study samples with heterogeneous early-life backgrounds across a wide age range. There are no datasets enriched for early adversity that are large enough to obtain robust estimates of effect sizes given the difficulty of recruiting these types of samples. As a result, there are no quantitative studies despite numerous qualitative reviews and theoretical papers (Bick and Nelson, 2016; Brito and Noble, 2014; Callaghan and Tottenham, 2016a; McLaughlin et al., 2019; Tottenham, 2020). Thus, meta-analytic techniques are the *only* available quantitative method to understand the developmental pathways by which different adversity exposures influence brain structure across the first two decades of life.

Adversity is a broad, umbrella term used throughout the literature to describe a wide range of experiences. Developmental perspectives on early-life adversity propose that children’s neurobehavioral systems adapt to the demands of their ecological contexts in a way that supports their own survival needs (Johnson et al., 2015). Different environmental features, therefore, are expected to have at least partially distinct effects on brain development due to developmental constraints on sources of input and neuroplasticity mechanisms (Gabard-Durnam and McLaughlin, 2020; Ho and King, 2021). This developmental approach differs from approaches that define adversity in terms of sociolegally-defined categories primarily created to implement public policy, such as maltreatment, foster care, and institutional care. As a result, these distinctions are unlikely to have biological relevance (Smith and Pollak, 2021). Allostatic load, defined as the phenotypic consequences of chronic activation in stress response systems, has been proposed as a common mechanism across all early-life adversity types (McEwen, 1998). As several groups have noted (Callaghan and Tottenham, 2016b; Ellis and Giudice, 2014; McLaughlin and Sheridan, 2016), however, the allostatic load model is limited by its emphasis on mature adult systems and assumption that all adverse experiences impact the brain in the same manner.

Whether early adversity is interpersonal or not is a prominent ecological characteristic with extensive evidence spanning decades for having robust, unique effects on neurobiology and behavior across species (French and Carp, 2016; Meaney, 2001; Patel et al., 2019; Sanchez et al., 2015; Sapolsky, 2021; Villalta et al., 2018). Characterizing adverse experiences as being interpersonal or non-interpersonal is commonplace in non-human animal models and adult clinical samples, but rarely has been applied to developmental work in early-life adversity. Humans are an altricial species that rely on caregivers for survival, allostasis, and regulation of affective systems during a protracted developmental period (Callaghan and Tottenham, 2016a). During infancy and early childhood, therefore, developmental outcomes depend on proximal, interpersonal interactions with caregivers to a much greater extent than distal environmental influences that affect the child indirectly such as family income or the community (Ho and King, 2021). Indeed, primate research finds that social relationships have an outsized interplay with survival (i.e., frontolimbic) circuitry relative to non-interpersonal aspects of the environment (Sanchez et al., 2015; Sapolsky, 2021). Adverse events that directly involve or impact interpersonal relationships (e.g., emotional/physical maltreatment) violate children’s most basic safety assumptions, thereby threatening children’s affective perceptions of felt security (Smith and Pollak, 2021). Work in trauma-exposed children supports this notion, showing that interpersonal traumas exhibit more pronounced effects than non-interpersonal traumas on emotion regulation difficulties (Villalta et al., 2018), behavior problems (Price et al., 2013), and stressor- and mood-related psychiatric symptoms (Ford et al., 2011). Interpersonal early adversities have been suggested to preferentially impact affective brain regions (Bick and Nelson, 2016; Callaghan and Tottenham, 2016b), likely because the neural regions supporting emotional processing, learning, and responding (i.e., amygdala, striatum, hippocampus) are modulated by caregiving cues and undergo rapid structural growth during early life when children are most dependent on close others. Consistent with this proposal, disruptions to the most salient early interpersonal relationship (i.e., the caregiving environment) during infancy and childhood have particularly profound consequences for changes in limbic brain structure across species (Sullivan, 2012). Interpersonal and non-interpersonal adversities likely have divergent influences on brain structure, with interpersonal early adversities targeting ‘emotional’ limbic neurobiology.

Poverty and experiences stemming from socioeconomic inequality are distal adverse environmental contexts that are experienced very differently than interpersonal early adversity by the developing child. Indeed, poverty may be considered orthogonal to interpersonal adversity, although they may correlate at the population level. Poverty exposure is measured by household and neighborhood indicators of socioeconomic disadvantage, with poverty involving lower levels of income, education, and employment, poorer housing quality, and more crime in the community (Brito and Noble, 2014). Socioeconomic disadvantage and other non-interpersonal adversities influence the child indirectly via a cascade of events (Brito and Noble, 2014; de Mendonça Filho et al., 2023). Childhood socioeconomic disadvantage overwhelmingly impacts the availability of material resources and access to cognitive enrichment opportunities, and therefore may be more likely to influence cortical regions central to language, executive functions, and sensorimotor processing (Brito and Noble, 2014). Although interpersonal early adversity and poverty are modestly correlated, poverty is distinct because poverty does not *directly* disrupt attachment processes or threaten a child’s fundamental safety and security (Amso and Lynn, 2017). Indeed, emerging evidence shows divergent effects of interpersonal early adversity and early socioeconomic disadvantage on limbic brain structure in adults (Lawson et al., 2017) and neuroendocrine and immune function in children (de Mendonça Filho et al., 2023). This notion is supported even in the context of childhood violence; caregiver-perpetrated violence exposure has been linked to amygdala sensitization and in turn externalizing problems, whereas violence exposure outside of the home (school, community) was associated with amygdala habituation with no link to externalizing problems (Stevens et al., 2021). These contrasting neural adaptations may reflect differences in the interpersonal nature of these adverse experiences; caregiver-perpetrated violence may give rise to hypervigilance for imminent threats in interpersonal interactions and result in amygdala hyperreactivity to emotional face stimuli, while amygdala habituation in children exposed to school or community violence may indicate learning differences for social cues that do not involve close relationships (avoidance, enhanced threat/safety discrimination). Conceptually and neurobiologically, socioeconomic disadvantage and its ecological concomitants can be considered as a form of non-interpersonal early adversity.

As Tottenham and Sheridan highlight (2010), understanding the effects of early adversity also depends highly on the timing of measurement. Timing is particularly important given that characterizing the developmental course of a phenotype is essential to providing deeper insight into the mechanisms that give rise to the adult form. For example, the adversity-induced acceleration hypothesis is one theoretical account that proposes differential associations between particular early adversities and specific brain structures across development (Callaghan and Tottenham, 2016b). Namely, caregiving adversity (the primary form of interpersonal adversity early in life) appears to accelerate (perhaps temporarily) the timing of the development of brain regions integral to emotional processing and learning to meet the immediate demands of functioning independently (potentially at an age earlier than ‘expected’) (Gee et al., 2013). Although this phenotype may be immediately beneficial early in life, it may have long-term consequences for neuroplasticity and emotional learning at later ages (Hanson and Nacewicz, 2021; VanTieghem et al., 2021). It has also been suggested that lower socioeconomic status environments may hasten the pace of neural development in children through broader domain-general neuroplasticity mechanisms involving the acceleration of biological aging across many bodily systems (Tooley et al., 2021). Alternative proposals posit that socioeconomic disadvantage is linked to cognitive and neurodevelopmental delays (Brito and Noble, 2014; Rakesh and Whittle, 2021). Characterizing the influence of early adversity exposure through the lens of development is critical for informing models of equifinality and multifinality that explain heterogeneity in adversity exposure types and brain volumetric outcomes.

Empirically measuring brain structure across the first two decades of life in a large study sample enriched for heterogeneous adversity types is currently a nearly impossible task. Therefore, this preregistered meta-analysis sought, for the first time, to quantitatively summarize the association between early-life adversity and brain structure throughout the early lifespan and investigate relevant moderating influences. Specifically, we hypothesized that age at measurement and adversity type (interpersonal versus socioeconomic disadvantage) would exhibit regionally-differential associations with structural development from infancy to adolescence. Understanding the multiple pathways by which adversity exposure influences brain structure is a critical step toward characterizing the mechanisms that may give rise to potentially domain-specific outcomes following early-life adversity exposure.

## 2. Results

### 2.1. Sample characteristics

The literature search identified 6,964 abstracts after removing duplicates (*n*=1,305). A total of 92 unique samples from 81 articles were included in this meta-analysis after eligibility screening (Figure 1; *Supplementary Information*, Table S1); 65% were characterized by interpersonal early adversity (*n*=60) and 35% by socioeconomic disadvantage (*n*=32). The included studies comprised 27,234 youth with mean sample ages between the ages of 1-month and 18 years (*M*=10.39 years). The average gender distribution across studies was 49% female and 51% male. Most samples were collected in North America (*n*=65, 70%), with the rest occurring in Asia (*n*=7, 8%), Australia (*n*=4, 4%), Europe (*n*=14, 15%), and South Africa (*n*=2, 2%). Among studies of interpersonal early adversity, 88% were clearly characterized by caregiving-related early adversities (*n*=53). The remaining 11% of studies of interpersonal early adversity (*n*=7) likely involved the caregiver (e.g., being hit by someone), but it was unclear who the perpetrator was from the original reporting. Some studies reported the timing of early-life adversity (*n*=24), with a mean age of onset of 22 months (*SD*=23) that ranged from birth to 5 years.

**Figure 1.**
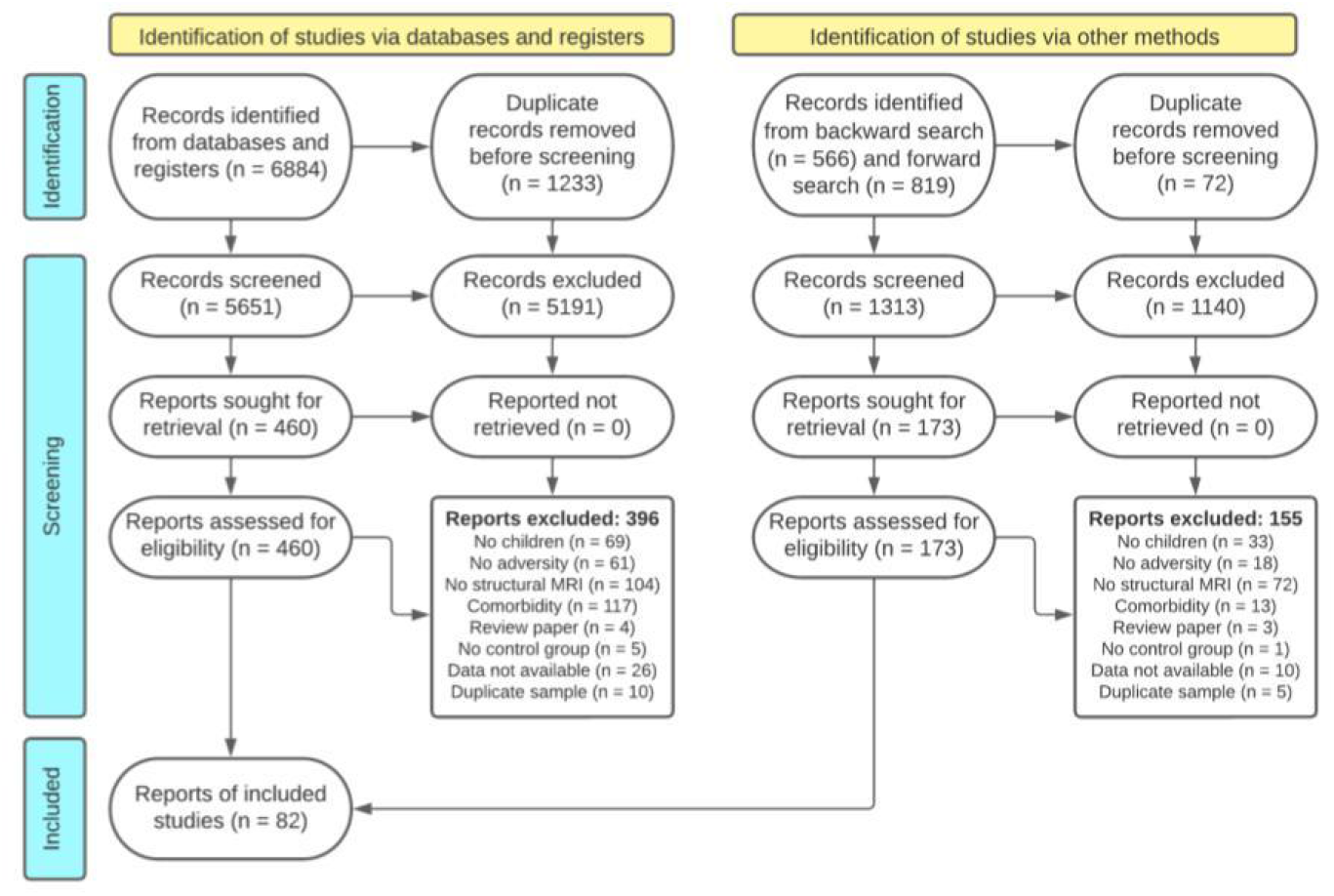
PRISMA flow diagram for systematic review.

### 2.2. Age-dependent effects of early-life adversity exposure (age treated as a variable of interest)

We observed divergent age-related associations for interpersonal early adversity and socioeconomic disadvantage with regard to specific brain volumes across childhood and adolescence (Figure 2; *Supplementary Information*, Table S2). Interpersonal early adversity exposure (vs. no exposure) was associated with larger amygdala, hippocampus, ventral anterior cingulate cortex (vACC), ventromedial prefrontal cortex (vmPFC), and ventrolateral prefrontal cortex (vlPFC) volumes when measured in childhood. There was a subsequent directional switch after 12-years-old; interpersonal early adversity (vs. no exposure) was associated with increasingly smaller volumes in these regions across adolescence. Early socioeconomic disadvantage showed different age-related patterns in temporal-limbic structures. Specifically, greater socioeconomic disadvantage (vs. less) was associated with smaller volumes in the amygdala, hippocampus, parahippocampus, and all temporal gyri across childhood (until age ~12). The magnitude of these negative associations became increasingly attenuated with age until there were negligible volumetric differences between youth with and without exposure to socioeconomic disadvantage in these temporal-limbic regions during middle-to-late adolescence. Considering age and adversity type accounted for, on average, 78% (*SD=31;* range=19-100%) of the between-study heterogeneity in these associations between early-life adversity and brain volume. In contrast to these findings that considered age, the divergent course of brain development was obscured when we examined the effects of early-life adversity type without a careful consideration of age (when age was a covariate; *Supplementary Information*, Table S2 and Figure S2).

**Figure 2.**
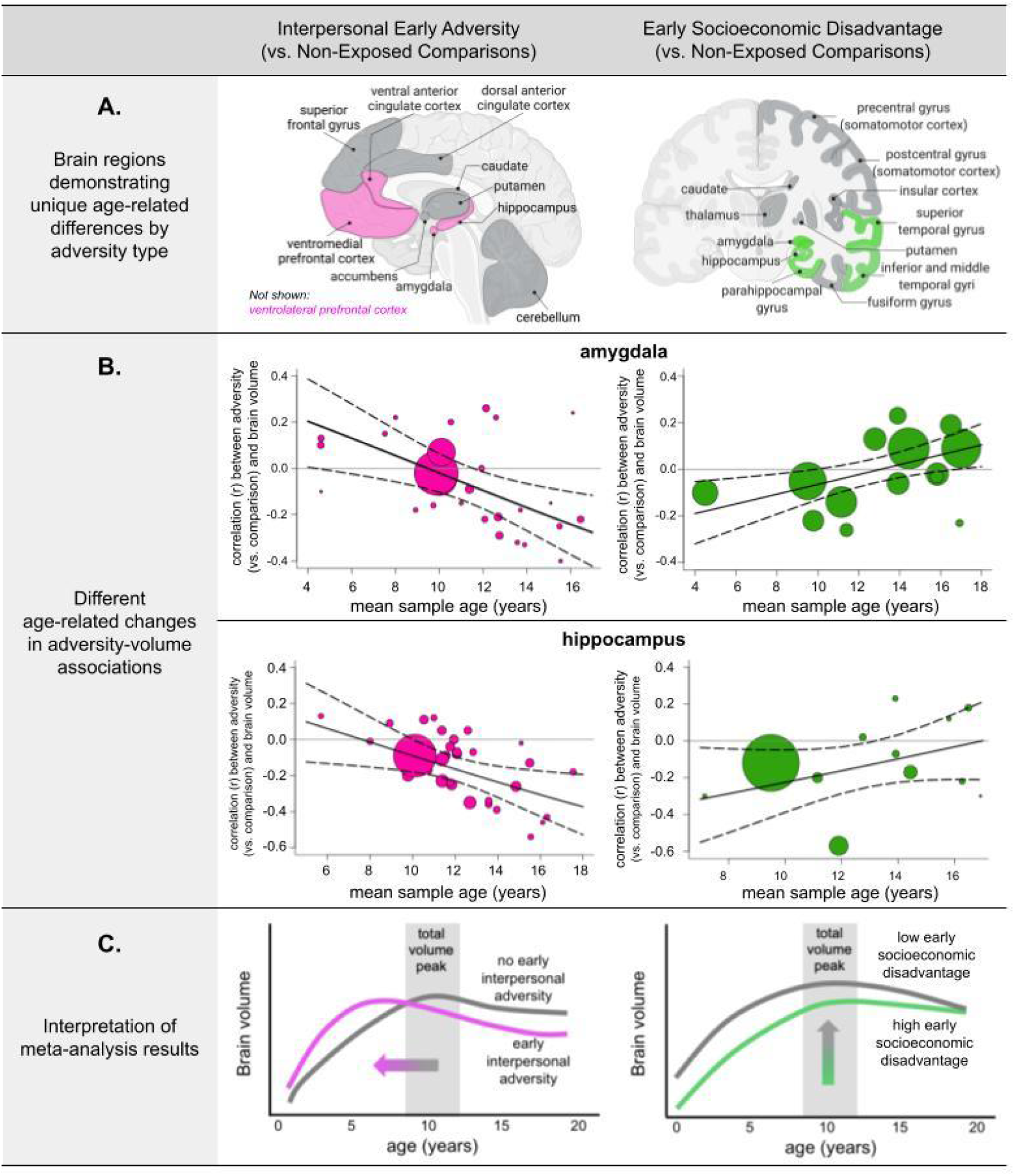
Age-dependent associations between early-life adversity and brain volume. **A.** All age-dependent associations indicate that the magnitude of the association between early adversity exposure (vs. no exposure) and brain volume changes with age. **pink** = <u>negative</u> age-related change with interpersonal early adversity (vs. non-exposed comparison); **green** = <u>positive</u> age-related change with higher (vs. lower) early socioeconomic disadvantage; **dark gray** = no age-related associations with either type of early adversity exposure. **B.** Amygdala and hippocampus show different age-related associations for interpersonal early adversity exposure (vs. no exposure) and early socioeconomic disadvantage. The size of the points reflects the relative sample size. Other regions not shown here exhibit comparable patterns of age-related change for interpersonal early adversity (ventral anterior cingulate cortex [vACC], ventrolateral prefrontal cortex [vlPFC], ventromedial prefrontal cortex [vmPFC]; see *Supplementary Information*, Figure S24) and early socioeconomic disadvantage (parahippocampus; inferior, middle, and superior temporal gyri; see *Supplementary Information*, Figure S25). **C.** Interpersonal early adversity preferentially shapes frontolimbic circuitry, with age-related changes supporting adversity-induced acceleration models. That is, interpersonal early adversity may contribute to early adversity-induced acceleration in childhood with a long-term tradeoff of later allostatic overload and attenuated neural plasticity. Early socioeconomic disadvantage preferentially shapes cortical-limbic structures.

### 2.3. Robustness of results

Leave-one-out analyses suggested that the findings reported here were robust. That is, the effect held even when removing samples where an average age at measurement had to represent a large age range. There was inconsistent evidence for publication bias, such that one metric suggested the presence of bias whereas the other metric indicated no bias, for the cerebellum, insula, inferior parietal cortex, inferior temporal gyrus, and occipital lobe (*Supplementary Information*, Table S2). Publication bias was not evident for other brain regions

### 2.4. Post-hoc exploratory analyses

Post-hoc exploratory interactions of early-life adversity-volume associations were examined for hypothesis generation and to facilitate considerations for future research.

#### 2.4.1. Methodological factors

Exploratory analyses evaluated the extent to which study methods were associated with changes in the magnitude of adversity-volume associations (*Supplementary Information*, Table S3). Methodological factors that could be coded across all studies included sex assigned at birth, total volume correction, scanner type, segmentation method, and magnetic field strength. Small, but negligible, study gender differences in adversity-volume associations were identified for 14 of the 22 brain regions examined. Methodological factors did not impact the strength of the associations between early-life adversity and brain volume for most brain regions. No systematic differences in effects of methodological factors were observed.

#### 2.4.2. Threat and deprivation exposures

Exploratory analyses were conducted with adversity type coded as threat or deprivation because this framework has empirical support and could be coded from the available data, albeit less clearly than interpersonal and socioeconomic disadvantage types (*Supplementary Information*, Table S4). Threat and deprivation could not serve as the primary adversity types of interest for this study because threat is coded primarily from maltreatment or trauma exposure (Johnson et al., 2021), which often collapses across threat (e.g., physical/sexual abuse) *and* deprivation (e.g., physical neglect) constructs in the extant literature. Only two brain regions (the amygdala and caudate) showed evidence of age-related differences as a function of adversity type (threat vs. deprivation). For both, threat exposure (vs. no exposure) was not associated with brain volume in children, but was associated with increasingly smaller volumes across adolescence relative to non-exposed comparisons. No age-related differences were found for links between deprivation (vs. no exposure) and brain volume.

## 3. Discussion

Conducting a preregistered meta-analysis with 92 unique samples comprising over 27,000 youth, here we demonstrated an interaction between developmental timing and adversity type for brain volume such that interpersonal early adversities and socioeconomic disadvantage were associated with distinct developmental correlates that differed as a function of brain region and age at measurement. Specifically, interpersonal early adversity exposure (vs. no exposure) was primarily associated with larger frontolimbic volumes early in life, but smaller volumes in adolescence. By contrast, socioeconomic disadvantage was associated with smaller temporal-limbic volumes early in life, but these associations became non-significant later in adolescence. Leave-one-out analyses indicated that the results were robust. These findings indicate that the links between early-life adversity exposure and neurodevelopment are highly dependent on the type of experience, the age at which brain volume is measured, and the region examined.

The human brain undergoes significant construction during the first two decades of life and remains in a state of constant, dynamic change across this period. Molecular and cellular mechanisms of neuroplasticity also shift throughout development (Gabard-Durnam and McLaughlin, 2020; Ho and King, 2021). It should therefore be unsurprising that adversity-related phenotypes would also demonstrate notable dynamics across this developmental period. Nonetheless, while there currently exist many theoretical perspectives regarding the ability of developing systems to adapt to adverse environments (e.g., Callaghan and Tottenham, 2016b; Ellis et al. 2022; Johnson et al. 2016; McLaughlin and Sheridan, 2016; Smith and Pollak, 2021), there remains a limited consideration of the influence of development itself on resulting outcomes. That is to say, extant assumptions that adversity exhibits ontogenetically uniform effects do not adequately account for the importance of the ongoing change inherent to the process of development. Case in point—without investigating the role of age at assessment, our results would have falsely indicated that there was no association between interpersonal early adversity and amygdala volume (*Supplementary Information*, Table S2). However, considering developmental timing revealed that following interpersonal adversity (vs. no exposure), the amygdala and other brain regions central to emotional processing like the hippocampus, vACC, vmPFC, and vlPFC, were larger in early-to-middle childhood, but in adolescence, these regions switched to being smaller.

These findings support the adversity-induced acceleration model posited by Callaghan and Tottenham (2016b), which hypothesizes that early-life adversities, most notably those that involve direct insults to the parent-child relationship, instantiate a neural activity-based acceleration observed early in development that may alter the developmental trajectory of a given phenotype. The interpersonal early adversities included in this meta-analysis (as well as most interpersonal adversities that occur at a young age) overwhelmingly involved the caregiver. According to the acceleration framework, neural alterations observed following interpersonal adverse experiences may be best understood as adaptations that occur in response to environmental signals that indicate protection from a primary caregiver is unavailable or unreliable (Callaghan and Tottenham, 2016b). Development of relevant neurocircuitry may therefore temporarily accelerate to aid in the young child’s independent survival (i.e., navigation of safety versus danger) (Callaghan and Tottenham, 2016b). This acceleration is hypothesized to be the consequence of neural activity-based processes occurring at too-young an age; for example, in circuits that would support independent navigation of potential dangers at an age earlier than typical. Neural accelerations following interpersonal early adversity, therefore, are expected to be region-specific (i.e., not global in the brain). The current results demonstrating brain region-specificity in the observed associations support this hypothesis and are consistent with experimental work. Early caregiving adversity in rodents initially induces the overexpression of brain-derived neurotrophic factor (BDNF) early in development through epigenetic mechanisms, which causally increases axonal branching and dendritic arborization (and therefore volume) focally in the amygdala, hippocampus, and prefrontal cortex (Bennett and Lagopoulos, 2014). Behaviorally, larger amygdala and hippocampal volumes are associated with enhanced threat/safety learning (McLaughlin et al., 2016), but also higher anxiety (French and Carp, 2016; Roth et al., 2018), in youth and juvenile primates exposed to interpersonal early adversity.

Precocious neuroaffective development may abbreviate plasticity in frontolimbic regions later in life due to the early closure of sensitive periods for emotional learning (Callaghan and Tottenham, 2016b), such that individuals exposed to interpersonal early adversity demonstrate smaller brain volumes later in childhood and adolescence, as observed in the current meta-analysis and prior longitudinal work (Luby et al., 2019; VanTieghem et al., 2021). Indeed, several groups (Hanson and Nacewicz, 2021; Teicher et al., 2016; Tottenham and Sheridan, 2010) have posited that early adversity-induced enlargements may sensitize the amygdala to future stressors, resulting in smaller amygdala volumes at later ages. In rodents, frontolimbic volume shrinkage and reduced plasticity at later ages is induced by the long-term depression of BDNF expression following interpersonal early adversity (Bath et al., 2013), implicating synaptic pruning and dendritic atrophy as potential mechanisms. Accelerated frontolimbic myelination may also account for earlier, but attenuated volumetric growth (Gur et al., 2019). During adolescence, smaller frontolimbic volumes linked to interpersonal adversity are associated with poorer emotional awareness (Luby et al., 2017), higher levels of depression (Weissman et al., 2020), and (for ventral prefrontal regions only) worse attention and externalizing problems (Gold et al., 2016; McLaughlin et al., 2014). Although speculative, the developmentally hierarchical relationship between the early-developing amygdala and the later-developing prefrontal cortex may underlie the observed cortical associations with interpersonal early adversity (Tottenham and Gabard-Durnam, 2017).

Unlike interpersonal early adversity, socioeconomic disadvantage was associated with smaller amygdala, hippocampal, parahippocampal, and temporal gyri volumes in childhood, but by middle-to-late adolescence, these associations were no longer observed (Figure 2). In this meta-analysis, socioeconomic disadvantage encompassed poverty and ecological concomitants (e.g., neighborhood deprivation and crime). These results are consistent with longitudinal work showing attenuated temporal-limbic growth throughout childhood (Barch et al., 2021; Whittle et al., 2017) and faster cortical thinning (Piccolo et al., 2016) in relation to socioeconomic disadvantage. Similar to prior work on cortical surface area (Noble et al., 2015), socioeconomic disadvantage (vs. no exposure) was also associated with smaller prefrontal and parietal structures in this meta-analysis despite no age-related differences. Day-to-day stress and caregiving (support/hostility) partially account for links between early socioeconomic disadvantage and brain structure alterations in children (Luby et al., 2013). However, these brain structure findings may differ dramatically from the interpersonal early adversity results because socioeconomic disadvantage is largely linked to events that do not directly interfere with the interpersonal/attachment and emotional needs of childhood; that is, it is non-interpersonal in nature.

Instead, the neurodevelopmental effects of early socioeconomic disadvantage might be best explained by a link between wealth advantages, like cognitive enrichment, diverse language exposure, and complex sensory stimulation, and lateral cortical structures (Amso and Lynn, 2017). Unmet material needs, toxins, or poor nutrition may also explain these observations of smaller brain structures linked to socioeconomic disadvantage (Johnson et al., 2016). These associations appear to equalize during adolescence, perhaps suggesting that these early experiences may no longer be associated with the brain once children gain greater independence over their environment (Ho and King, 2021). Although the mechanisms remain elusive, curtailed synaptogenesis and cortical gyrification and/or a faster pace of synaptic pruning and myelination may play a role in the observed age-dependent associations between socioeconomic disadvantage and brain volume (Brito and Noble, 2014; Tooley et al., 2021). The role of biological aging mechanisms are inconsistent (Bush et al., 2018; Gur et al., 2019), perhaps in part due to unmeasured, co-occuring interpersonal adversities. Nonetheless, the observed associations between socioeconomic disadvantage and brain structure have functional relevance, with links to higher levels of depressive symptoms (Barch et al., 2020) and poorer language skills, cognitive control, and working memory (Taylor et al., 2020). These neural adaptations also may contribute to skills for navigating the ecological demands of impoverished environments, such as enhanced cognitive flexibility, persistence in obtaining immediate rewards, and faster detection of changing action-outcome contingencies (Frankenhuis and Nettle, 2020).

Though categorizing adversities based on their interpersonal involvement was theoretically motivated, there were alternative ways to parse adverse experiences (Ellis and Giudice, 2014; McLaughlin and Sheridan, 2016; Smith and Pollak, 2021). Another codable distinction was categorizing experiences by threat versus deprivation, a method of division supported by a growing body of work (McLaughlin et al., 2019). Therefore, threat versus deprivation was an important adversity type distinction to test in the current meta-analysis. These analyses indicated smaller overall volumes associated with early threat, deprivation, or both exposure types in most brain regions that did not change with age (i.e., age-invariant), except for the amygdala and caudate, which demonstrated no effects during childhood but reduced volumes in adolescence. Both approaches accounted for a large proportion of the variance in between-study heterogeneity. However, a greater proportion of variance was explained when dividing adversities by their interpersonal nature relative to dividing adversity based on threat versus deprivation. Consistent with a wealth of early-life adversity studies in nonhuman animals (French and Carp, 2016; Meaney, 2001; Patel et al., 2019; Sanchez et al., 2015; Sapolsky, 2021; Villalta et al., 2018), these findings suggest that interpersonal factors are critical to developmental outcomes. Importantly, there are shared features between socioeconomic disadvantage and deprivation distinctions. For example, operationalizations of deprivation overwhelmingly include the experience of poverty (Ellis et al., 2022). In contrast, the current meta-analysis differed from a threat versus deprivation approach with regard to how it treated emotional neglect/deprivation. Categorizing adversities by their interpersonal nature considered emotional neglect as more functionally similar to other forms of parental maltreatment, including abuse, than to poverty, whereas the threat versus deprivation distinction considers emotional neglect and poverty (and related exposures such as cognitive and material deprivation) as more mechanistically alike. The current meta-analysis provided empirical support for an interpersonal division by indicating that for very young children, emotional neglect/deprivation has different neurodevelopmental sequelae than impoverished financial environments, and may operate more like other forms of maltreatment (e.g., abuse). The extant literature is mixed on this point, so additional empirical tests of this hypothesis are warranted.

Findings from this meta-analysis underscore that the careful consideration of development facilitates a deeper appreciation of how the developing brain adapts to specific experience types. There are nonetheless important limitations to acknowledge when making inferences from this meta-analysis. There was notable heterogeneity in the methods used across studies to capture adversity due to the secondary data analysis. This precluded the systematic investigation of additional neurobiologically-relevant aspects of the adverse environments such as the timing of adversity exposure (Tottenham and Sheridan, 2010), chronicity, and environmental predictability (Cohodes et al., 2021). However, using an easily distinguishable characteristic, like whether or not the adverse experience directly involved disruptions to close interpersonal-affective relationships, did enable us to reduce heterogeneity and facilitate orthogonalization more easily than with other methods of adversity categorization. There was data sparsity in infant and toddler-aged samples, which may reflect numerous challenges involved in collecting MRI data at very young ages, especially in the context of adversity exposure. Further, though we attempted to capture age-related effects, we investigated associations cross-sectionally, precluding inferences about directionality. The mean sample age at scanning was also a relatively coarse measure given that the included samples often encompassed a wide age range. Leave-one-out analyses suggested that the inclusion of these samples did not bias the results. Nonetheless, long-term longitudinal work will provide a more accurate account of associations between adversity and age-related change in brain structure (VanTieghem et al., 2021). The articles included in this meta-analysis may have focused on one specific adversity type (e.g., violence), but the participants in the studies may have nonetheless been exposed to other unmeasured adversities. Results were robust to the influence of outlier studies, but there was some evidence of publication bias for the cerebellum, insula, inferior parietal cortex, inferior temporal gyrus, and occipital lobe. As such, the magnitude of early-life adversity-volume associations may be inflated in these regions. Brain region-of-interest (ROI) random-effects meta-analyses utilize optimal statistical methods for estimating effect sizes and for assessing reliability and between-study heterogeneity (Radua and Mataix-Cols, 2012). However, the included studies often have biased and limited inclusion of ROIs (Radua and Mataix-Cols, 2012), which was evident in the abundance of studies focused on the amygdala and hippocampus relative to other regions. Nonetheless, the results of this meta-analysis generated compelling hypotheses that should be replicated and expanded upon with whole-brain studies of early-life adversity and a broader range of brain structure measures.

Meta-analytic techniques allow for a more robust characterization of the association between early adverse experiences and the developing brain than is presently possible in the empirical literature. Though existing reviews have thoroughly summarized the link between early-life adversity and brain structure qualitatively (Bick and Nelson, 2016; Callaghan and Tottenham, 2016a; McLaughlin et al., 2019; Tottenham, 2020), few meta-analyses have quantitatively summarized these associations. Those that do exist largely have focused on adult populations.(McLaughlin et al., 2019) Only one meta-analysis on adversity and brain structure has included children to our knowledge (Lim et al., 2014), although the combined sample size of children and adolescents was much smaller than that of the adults (*n*=56 vs. *n*=275). By gathering the largest neuroimaging sample of adversity-exposed youth to date, the current meta-analysis demonstrates that adversity does not have a uniform impact on the developing brain, but instead displays age-, region- and experience-specific developmental associations. The current work proposes novel hypotheses about the mechanisms by which adversities might impact brain structure, serving as a foundation to inspire further investigation into important questions concerning the lasting influence of early adversity on development.

## 4. Materials and Methods

### 4.1. Study Inclusion and Exclusion Criteria

To be eligible for inclusion in the meta-analysis, studies need to have: (1) a human sample between the ages of birth and 18-years-old; (2) a measure of postnatal early-life adversity that occurred between prior to age 18-year-old, including exposure to child maltreatment, caregiving disruptions, caregiver psychopathology, low family or neighborhood socioeconomic status, and/or neighborhood crime, violence or lack of safety; (3) a measure of brain volume derived from magnetic resonance imaging (MRI) methods (T1-weighted MPRAGE); and (4) sufficient quantitative information to calculate at least one effect size for the association between early-life adversity and brain volume. Studies were excluded if they did not meet all four inclusion criteria.

### 4.2. Literature Search

This review adhered to the guidelines described in the Preferred Reporting Items for Systematic Reviews and Meta-Analyses (PRISMA) statement (Page et al., 2021). PubMed, PsycInfo, Medline, and Proquest Dissertations and Theses were searched to identify potential studies on January 22nd, 2021 and again on January 14, 2022. The following search terms were entered into each database to search all fields: (“infan*” OR “child*” OR “adolesc*” OR “toddler” OR “teen” OR “preschool*” OR “youth” OR “develop*”) AND (“child* advers*” OR “child* maltreatment” OR “child* abuse” OR “child* neglect” OR “maltreat*” OR “abuse*” OR “adverse child* experience*” OR “early life stress*” OR “advers* child* exp*” OR “child* trauma” OR “foster care” OR “kinship care” OR “institutional care” OR “disrupted caregiving” OR “institutional* youth” OR “social status” OR “poverty” OR “socioeconomic status” OR “socioeconomic disadvantage” OR “child* violence” OR “child* deprivation” OR “parent* addict*” OR “parent* depress*” OR “maternal depress*” OR “maternal addict*” OR “parent* psychopathology” OR “parent* mental illness” OR “parent* separat*” OR “domestic violence”) AND (“brain volume” OR “brain structure” OR “structural magnetic resonance imaging” OR “MRI” OR “neuroimaging” OR “voxel-based morphometry” OR “brain develop*” OR “neural develop*”). To ensure saturation, existing reviews and eligible studies were used to identify additional studies by inspecting the reference sections (“backward” search) and results of the “cited by” function in GoogleScholar (“forward” search). Citations and abstracts were exported as an RIS file and uploaded to Rayyan (Ouzzani et al., 2016), a web-based application to assist with systematic reviews, for screening.

### 4.3. Systematic Review and Study Selection

Trained research assistants independently screened the study abstracts to identify potentially eligible studies. Subsequently, the co-first authors (AF, AV) independently reviewed full-text manuscripts for determination of study inclusion. There was high interrater agreement regarding decisions to include or exclude studies at the abstract screening stage (*κ=.71*) and full-text review (*κ*=.92). The co-first authors (AF, AV) and senior author (NT) resolved decision disagreements through discussion of the study selection criteria.

### 4.4. Data Extraction and Management

Data extraction was performed using a standardized spreadsheet with the following columns: author(s), publication year, journal, sample size, mean sample age at MRI, sample gender breakdown, early-life adversity measure, brain region, scanner type, segmentation method, scanner magnetic field strength, and the statistics for the association between early-life adversity and brain volume. Two research assistants coded studies independently, and discrepancies were resolved through group discussion with the co-first authors (AF, AV). Interpersonal early adversity was coded for measures of caregiving disruptions (*n*=15; institutional care, caregiving instability, caregiver separation), caregiver psychopathology (*n*=12; anxiety, depression, addiction), maltreatment (*n*=23; emotional abuse, emotional neglect, physical abuse, physical neglect, sexual abuse), and interpersonal trauma (*n*=10; defined by an overall trauma score that included adversity exposures related to caregiving disruptions, maltreatment including abuse or neglect, and domestic violence exposure). At least one primary caregiver was, by definition, involved in the vast majority of interpersonal early adversities. Socioeconomic disadvantage was coded for indicators of low family socioeconomic status (*n*=25; income, income-to-poverty/needs ratios, poverty status, parental education), non-interpersonal trauma (*n*=1), community crime/violence exposure (*n*=2), neighborhood poverty (*n*=3), and area deprivation (*n*=1).

### 4.5. Data Analysis

A minimum of seven studies was required to ensure sufficient power (Jackson and Turner, 2017), and thus meta-analyses were conducted for 22 brain regions (*Supplementary Information*, Figure S1).

#### 4.5.1. Effect size calculations

Pearson’s correlation coefficients (*r*) were the effect size indices used to quantify the magnitude of the associations between early-life adversity and brain volume. When studies reported other effect size measures, the effects were converted to Pearson’s correlation coefficients using standard methods (Lenhard and Lenhard, 2016). Only one effect size per unique sample, per brain region was used in the meta-analysis to ensure independence.

#### 4.5.2. Combining and comparing effects across studies

An inverse variance-weighted, random-effects meta-analysis was conducted for each brain region using the *metafor* 2.4-0 package (Viechtbauer, 2010) in *R-4.0.2* (R Core Team, 2020). Random-effects meta-analyses estimate the fixed, or average, effects as well as between-study heterogeneity in the magnitude of the associations between early-life adversity exposure and brain volume. Random effects meta-analyses assume that there are inherent methodological differences across studies (e.g., sampling, scanner). Moreover, random-effects models facilitate the examination of systematic, theoretically-driven sources of heterogeneity that account for the between-study variability in effect sizes beyond these inherent study differences. The correlation coefficients were converted to the Fisher’s *z* scale to obtain a normal sampling distribution. Weighted effect sizes with confidence intervals that did not include zero were considered meaningful.

#### 4.5.3. Heterogeneity

Heterogeneity of effect sizes across studies was evaluated using the Chi-square *Q* statistic and the *I^2^* index (Higgins and Thompson, 2002). Mixed-effect meta-regression models with estimation via full information maximum likelihood were conducted to investigate factors that may account for between-study heterogeneity. Independent variables included age at MRI scan (sample mean), adversity type (1=interpersonal vs. 0=socioeconomic disadvantage), age x adversity type interaction, sample distribution of sex assigned at birth (% female), and total volume correction (1=yes vs. 0=no).

#### 4.5.4. Publication bias

The presence of publication bias was assessed with Egger’s test of funnel plot asymmetry (Egger et al., 1997) and the trim-and-fill method (Duval and Tweedie, 2000). Funnel plots depicted study effect sizes (x-axis) as a function of their standard errors (a reflection of sample size; y-axis), which appear asymmetrical in the presence of publication bias as a result of smaller samples reporting larger effect sizes. Publication bias was considered evident when *p*<.05 for Egger’s test, indicating that the funnel plot was asymmetric, and when the trim-and-fill method estimated that one or more studies needed to be imputed to make the funnel plot symmetrical.

#### 4.5.5. Model robustness

Leave-one-out analyses assessed the robustness of all results.

#### 4.5.6. Deviations from preregistration plan

This preregistered study intended to conduct brain region-of-interest and whole-brain coordinate-based meta-analyses given that they possess different advantages and limitations (Radua and Mataix-Cols, 2012). However, there were insufficient voxel-based morphometry studies (*k*=13) for a coordinate-based meta-analysis.

### 4.6. Data and Code Availability

The pregregistration plan, study data, analysis scripts, result outputs, and supplementary information supporting the findings of this study will available on the Open Science Foundation project for this study upon being accepted for publication.

## Acknowledgments

This work was supported by the National Institute of Mental Health (R01MH091864, R01MH091864-10S1; MPI: Tottenham and Milham). This funding source had no involvement in the meta-analysis design or analysis, writing the manuscript, or decision to submit the article for publication. Our PRISMA flow diagram was created with Lucidchart.com and our brain figures were created with BioRender.com. The following co-authors are now at other institutions: Ariel Katz at Yeshiva University; Eleanor Hansen at the National Institute of Mental Health; Nathan Martin at Palo Alto University and Stanford University School of Medicine; and Ayumi Tachida at the University of Washington School of Medicine.

